# Neuronal Branching is Increasingly Asymmetric Near Synapses, Potentially Enabling Plasticity While Minimizing Energy Dissipation and Conduction Time

**DOI:** 10.1101/2023.05.20.541591

**Authors:** Paheli Desai-Chowdhry, Alexander B. Brummer, Samhita Mallavarapu, Van M Savage

## Abstract

Neurons’ primary function is to encode and transmit information in the brain and body. The branching architecture of axons and dendrites must compute, respond, and make decisions while obeying the rules of the substrate in which they are enmeshed. Thus, it is important to delineate and understand the principles that govern these branching patterns. Here, we present evidence that asymmetric branching is a key factor in understanding the functional properties of neurons. First, we derive novel predictions for asymmetric scaling exponents that encapsulate branching architecture associated with crucial principles such as conduction time, power minimization, and material costs. We compare our predictions with extensive data extracted from images to associate specific principles with specific biophysical functions and cell types. Notably, we find that asymmetric branching models lead to predictions and empirical findings that correspond to different weightings of the importance of maximum, minimum, or total path lengths from the soma to the synapses. These different path lengths quantitatively and qualitatively affect energy, time, and materials. Moreover, we generally observe that higher degrees of asymmetric branching— potentially arising from extrinsic environmental cues and synaptic plasticity in response to activity— occur closer to the tips than the soma (cell body).

## 1 Introduction

The concept of asymmetry lies at the core of many biological processes, particularly in the nervous system, from asymmetries at the molecular level to whole-brain asymmetries. At the molecular level, asymmetry underlies the electrical and chemical transmission that enables information processing in the brain. Neurons connect to one another through axons and dendrites at synapses, where inter-cellular channels allow the transmission of signaling molecules, or neurotransmitters, and the spread of electrical currents. The asymmetry of these channels at the molecular level leads to functional asymmetry of the synapses, which is a key property enabling sensory processes [1]. At the cellular level, polarity and the asymmetric organization of cellular component is vital to many processes such as cell migration, cell division, and morphogenisis [3]. Asymmetry in neurons in particular has an important role in determining the physiology of neural circuits and cognition [4].

At the whole brain level, a key feature and an important topic in the study of the human brain is its division into hemispheres. The asymmetry between the left and right specialized regions of the human brain is crucial in our understanding of its structural organization and cognitive functions; many cognitive and psychiatric disorders are linked to specific alterations in this lateral hemispheric asymmetry [5]. Hemispheric asymmetries have been observed not only in humans, but in a range of species– including mammals, birds, reptiles, and fish–suggesting that lateral asymmetry is not unique to humans but rather an important principle in the structure and function of the nervous system [6].

In order to begin to understand broad-level asymmetries in the human brain, it is important to begin with the basic building blocks of the nervous system: neurons [7]. Neurons are said to be one of the most polarized cells in the body, with two distinct structural and functional domains— axons and dendrites [8]. A deeper understanding of the details of the structure and function of these neurites and how they respond to developmental and environmental cues to form synaptic connections is a crucial step leading up to an understanding of whole-brain asymmetry, cognition, behavior, and how alterations lead to diseased states [7].

Axons and dendrites form extensive branching trees that allow them to connect to one another, enabling information processing and communication in animals. Axons and dendrites are morphologically and functionally distinct; axons have long parent branches that can transmit information across large distances, and dendrites have shorter branches with more extensive branching trees. Axons utilize action potentials to transmit information over long distances, sometimes even crossing brain regions. The branching patterns and asymmetries of axons are characterized by systematic changes in branching radius and length across bifurcation branching points and are known to play a key role in signal propagation dynamics in neurons [9]. These axons connect to the dendrites of other neurons, which in contrast, rely on passive electronic spread and do not conduct action potentials [10]. Axons and dendrites have different mechanisms for forming new branches near their synaptic connections, allowing them to form the circuitry that is the backbone of information flow in the nervous system in the most efficient and frugal way [11, 12]. Foundational work by Santiago Ramón y Cajal documented the vast diversity of structural forms in neurons through detailed drawings of the morphology of neurons across cell-types. Ramón y Cajal established the correspondence between these diverse morphological forms and the vast functional diversity across cell-types by proposing functional principles that govern the structure such as conservation of space, time, and materials [13].

Previous work has attempted to develop a quantitative formalism to describe neurite branching through the laws of conservation of time and materials as described by Ramón y Cajal, using principles of optimization and a graph theoretical algorithm to generate biologically realistic synthetic axonal and dendritic trees [14, 15]. While this framework is able to successfully generate biologically accurate branching trees, it is limited in that it only considers the lengths of branching processes. Focusing on the 1-dimensional trace of these structures only captures one element of the biological factors that affect information processing speed, thus ignoring other important contributors. Foundational work by Hodgkin and Rushton describes the theoretical and empirical foundation for a quantitative description of the dependence of conduction velocity on the caliber of neurites as well as myelination [16, 17]. Our previous work incorporates volumetric interpretations of conduction time delay and material costs using mathematical principles from metabolic scaling theory in relation to cardiovascular networks to incorporate metabolic costs. Synthesizing these ideas leads to a unifying model that can predict various morphological structural parameters for axons and dendrites across a range of cell types [18].

We observe significant deviations from symmetric branching in neuron morphology data, as previewed in Figure 1D, suggesting that asymmetric branching is an important feature for the structure of neurons, likely corresponding to functional consequences as well. Although foundational work in modeling cardiovascular networks assumes that the branching junctions have perfect symmetry of the two daughter branches [19, 20], in biological resource distribution networks there is substantial variation around this symmetric case. Zamir first quantified deviations around symmetric branching that occur in vessels, showing differing levels of asymmetry across levels of coronary arteries [21, 22]. Further work by Tekin et al. built on this to establish systematic patterns in asymmetry throughout cardiovascular networks, adding to the analysis of asymmetry in branching length and width to incorporate patterns of asymmetry in branching angles [23], as well as deriving optimization principles that underlie these patterns.

**Fig 1.**
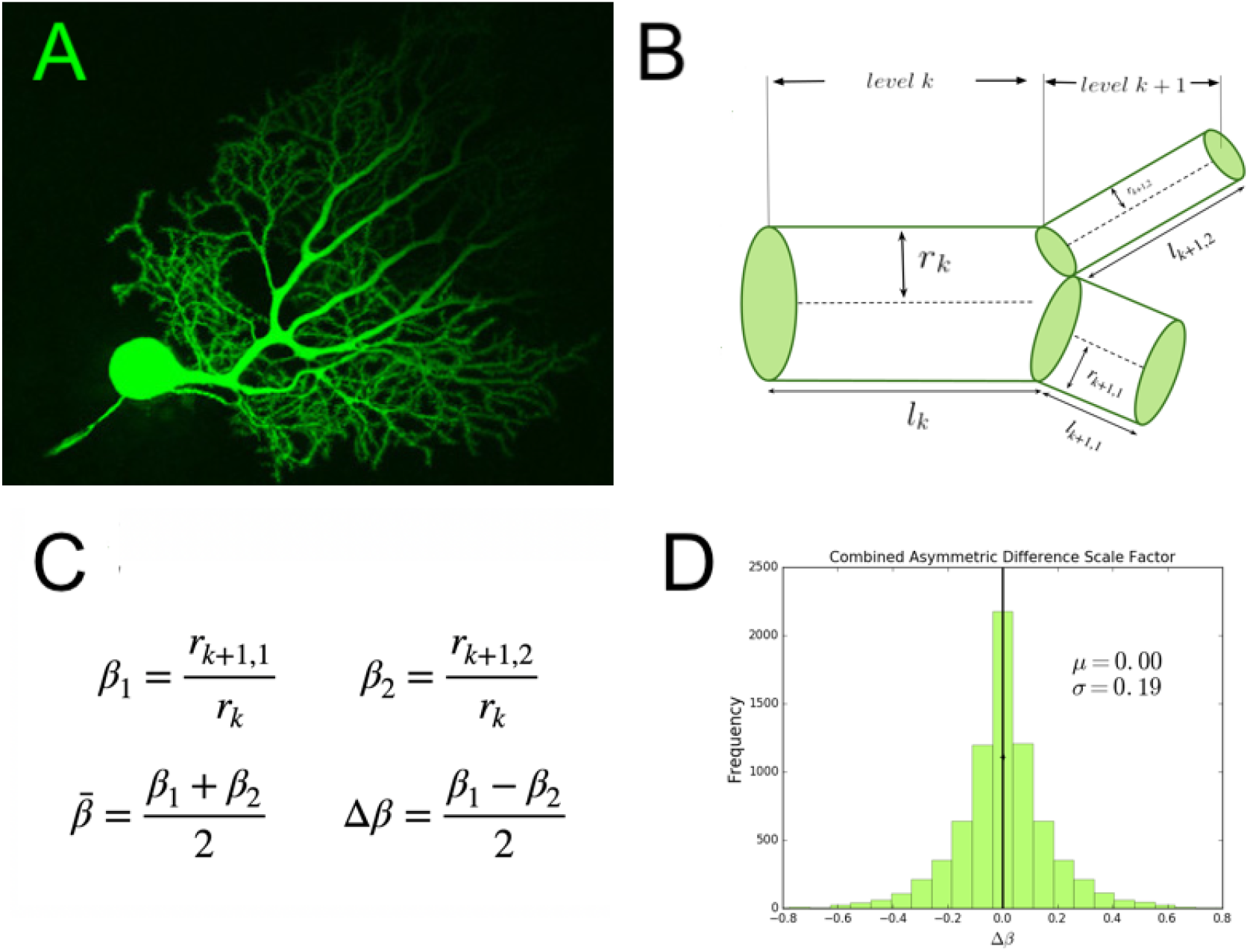
(A) An image of a mouse cerebellar Purkinje neuron and its dendritic branching structure. This image was obtained using confocal microscopy and Lucifer yellow fluorescent dye. We have cropped this image available on CellImageLibrary.Org, distributed by Maryann Martone, Diana Price, and Andrea Thor [2] (B) A diagram of a branching junction as part of a hierarchical branching network with successive branching levels, illustrating asymmetric branching junctions (C) Definitions of asymmetric scale factors, *β*_1_ and *β*_2_, and average and difference scale factors, 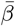 and *Δ β*, (D) A quantification of the branching asymmetry present across all data analyzed, as measured by the difference scale factor, *Δβ*, where the most symmetric values lie at a value of 0.

In order to understand the role of asymmetric branching in neuronal function across cell-types, here we extend our model of the structure-function correspondence to incorporate asymmetric branching. Using the asymmetric branching approach to model neurons, we must consider a multitude of path lengths from the soma to the synapses, suggesting that the whole network–rather than one optimal path–has an important contribution to neuron function and computation. Our results allow us to formulate hypotheses about the connection between branching and plasticity. In particular, our results suggest that it is possible that asymmetric branching emerges due to plasticity and responses to external factor. We hypothesize that asymmetric branching provides these dynamic branching processes with a robust architecture that is resilient to damage and allows them to adapt to fluctuating environments.

## 2 Theory

We represent neurons as hierarchically branching information processing networks, with successive branching levels that decrease in radius and length according to a scaling (i.e., power law) relationship. Figure 1 illustrates this with a representative image and a diagram of a branching junction.

We predict how the information processing function and surrounding substrate govern the branching structure of neurons. We do this by optimizing a mathematical cost function subject to a set of constraints, which allows us to obtain theoretical predictions for structural parameters that are the best possible given the biological constraints of the physical system [24]. Here, we choose a cost function that minimizes conduction time delay and energy consumption (represented by power loss) that is subject to computational, biological, and physical constraints.

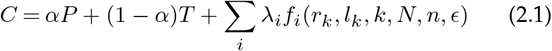

Equation 2.1 is a general form of this equation, where *T* is conduction time delay and *P* is power loss due to dissipation based on the assumption that these neuron processes are like wires through which a current is flowing, subject to electrical ohmic resistance. The parameter *α* can be toggled between 0 and 1 to minimize either power or time alone. The remaining terms in this function are constraint functions, representing biological quantities such as material costs that are held constant during the optimization. Each term in the cost function depends on the radius and length of the branch at each branching generation *k*, where 0 is the branching generation at the parent branch connected directly with the soma, and *N* is the last branching generation at the tips. The constraint functions *f*_*i*_ depend on the radius and length, *r*_*k*_ and *l*_*k*_, the branching ratio *n* (where n = 2 for a bifurcating function), and a parameter describing myelination, *ϵ*, where *ϵ=0* for unmyelinated fibers and 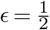 for myelinated fibers. We chose this parameter to vary this way because of previous foundational experimental and theoretical work that shows the conduction velocity is proportional to the square root of the diameter of a neuron fiber for unmyelinated processes and directly proportional to the diameter for myelinated fibers [16, 17]. Here, we focus on two main constraints: a material constraint, which we represent as the total network volume, and a time delay constraint, which we consider for the specific cases that focus on power minimization.

In our previous work, we use optimization methods to solve for theoretical predictions for scaling ratios for radius and length of processes in successive branching generations, 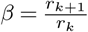 and 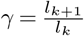 [18]. However, a key assumption of this work is that the branches are symmetric—the radius and length of the two daughter branches at each branching junction are identical. Despite this assumption, most biological axons and dendrites exhibit asymmetric branching [9, 25]. By analyzing neuron image reconstruction data from NeuroMorpho.Org [26], we quantify the pervasiveness of asymmetric branching across different cell types, as shown in Figure 1D.

In Figure 1B, we show an example of asymmetric branching. Here, we have two unequal daughter branches at the bifurcation point, so there are two separate scaling ratios for radius and length, 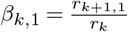 and 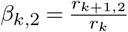(shown in Figure 1C), and 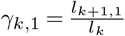 and 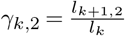 respectively.

We define the average scale factor as 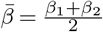 and the difference scale factor as 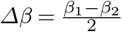 (shown in Figure 1C) based on conventions in previous work [27]. If we define *β*_1_ as the scaling ratio corresponding to the larger branch, we can describe *β*_1_ and *β*_2_ in terms of the average and absolute value difference scale factors as in Equation 2.2.

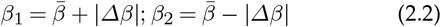

Thus, we can think of |*Δβ*| as a measure of the magnitude of the asymmetry, or the amount of shift away from the average. Figure 1D shows a distribution of *Δβ* in combined data for a range of cell types and species, preserving the sign as well as the magnitude to show variance around the symmetric case in both directions. We later break this data down into specific cell and process types in Figure 3.

Using an existing mathematical framework for asymmetric branching networks in the cardiovascular system [27], we extend our previous model [18] and are able to relax the assumption of symmetric branching. Using the scaling ratios in our expressions for power, time, and network volume along with the values for the radius and length at the tips, we derive whole network properties. As compared with our previous work, we needed to develop much more clever mathematical methods and do much more extensive derivations than for the symmetric theory. A big advance in overcoming these challenges is that we solve these equations recursively (See Appendix).

First, we define power, one of the functions to be minimized in the optimization, in terms of the asymmetric scale factors.

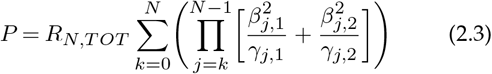

Note that we can also formulate this in terms of the difference and absolute value difference scale factors, where 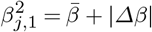 and 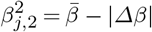. More explicit descriptions of the functions in this form can be found in the Appendix.

The other function to be minimized in the optimization is the conduction time delay. This term is more complicated with asymmetric branching, as there are multiple possible paths that a signal might take through the network. Previous work on plant networks deals with deviations from symmetry using a combination of terms relating to the mean and maximum path lengths [28]. Thus, we consider different cases of time delay: average time, total time, maximum time, and minimum time.

We define total time delay as follows in Equation 2.4.

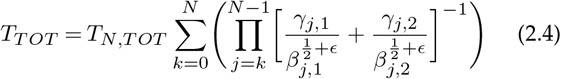

The average time delay is similar, though the total time at each generation is divided by the number of branches at that generation.

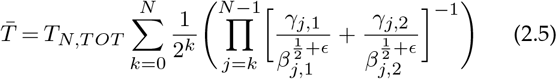

Finally, we define the time delay for the maximum and minimum path length. If we choose *r*_*k*+1,1_to be the larger daughter radius (Figure 1B), then we can define the maximum path length.

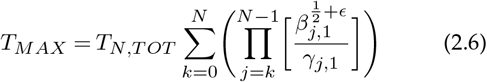

Similarly, we can define the minimum path length.

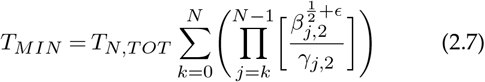

Next, we define the network volume or material cost— one of the constraint functions to be held fixed in the optimization.

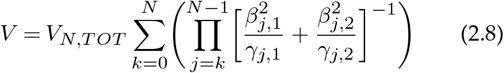

In this study, we will minimize the cost function in equation 2.1 under different limits to arrive at a suite of relationships between the two scaling ratios, *β*_1_ and *β*_2_, based on a scaling exponent P that dictates a generalized conservation equation:

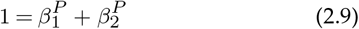

Although the distribution of scaling exponents yields information about broad network behavior, we first focus on how asymmetry changes with the distance from the soma. If we find that the asymmetry is localized to specific parts of the cell, this could be due to differences in the functional underpinnings that drive structures in different regions of the cell or due to other extrinsic factors such as connecting neurons or due to environmental cues. In order to analyze the data in terms of distance from the soma, we can use an established measure called leaf number that has been used to study scaling in dendritic branching [29]. The leaf number is defined as the number of tips that are distal to each branch. The leaf number at the tips will be equal to 0, and the leaf number will be greatest near the soma. Figure 2 illustrates leaf numbering. For each pair of radius scaling ratios in the data, we have a corresponding leaf number of the parent branch of the junction. We can define asymmetry level in terms of the difference between *β*_1_ and *β*_2_, or the difference scale factor in Equation 2.2. Distributions of this difference scale factor for different cell types are shown in Figure 3. Looking at the relationship between leaf number and this measure of asymmetry will allow us to determine where the most asymmetry occurs in terms of distance from the soma, as illustrated in Figure 4.

**Fig 2.**
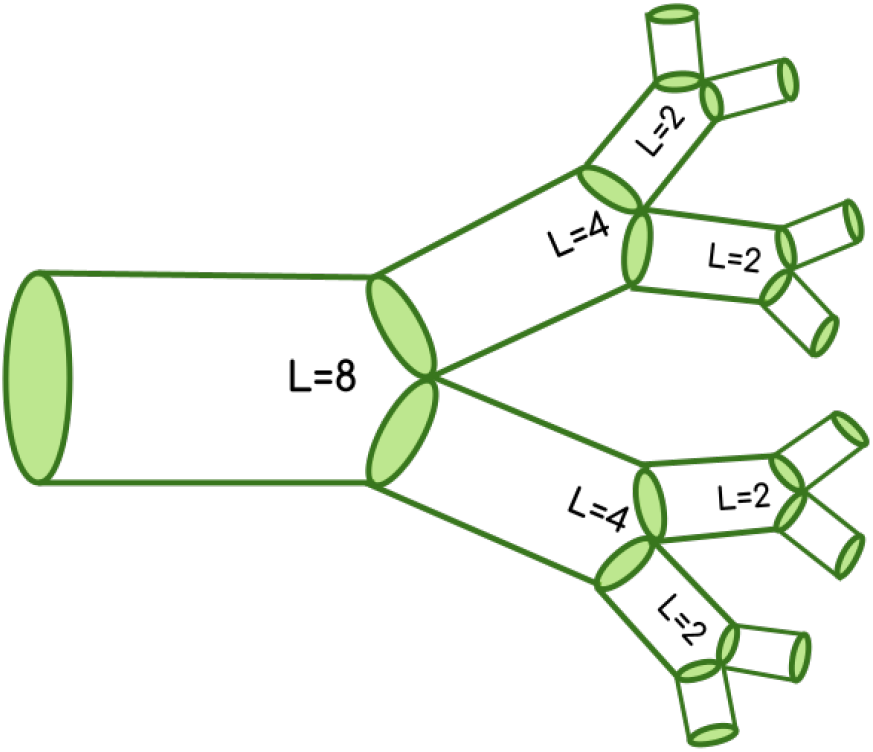
Branching Network with Leaf Number

**Fig 3.**
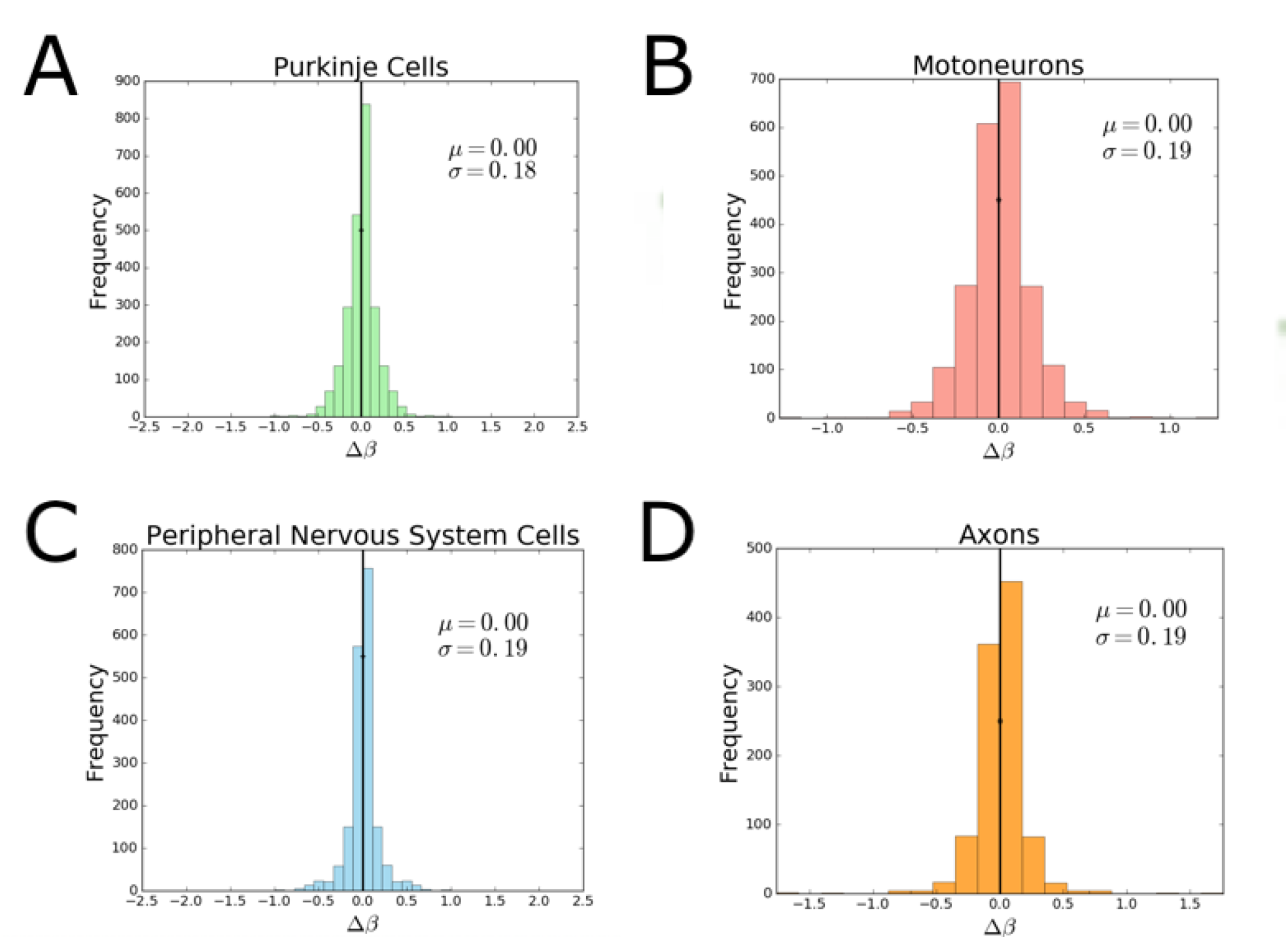
Plots of Asymmetric Difference Scale Factor for (A) Purkinje Cells, (B) Motoneurons, (C) Peripheral Nervous System Cells, and (D) Axons

**Fig 4.**
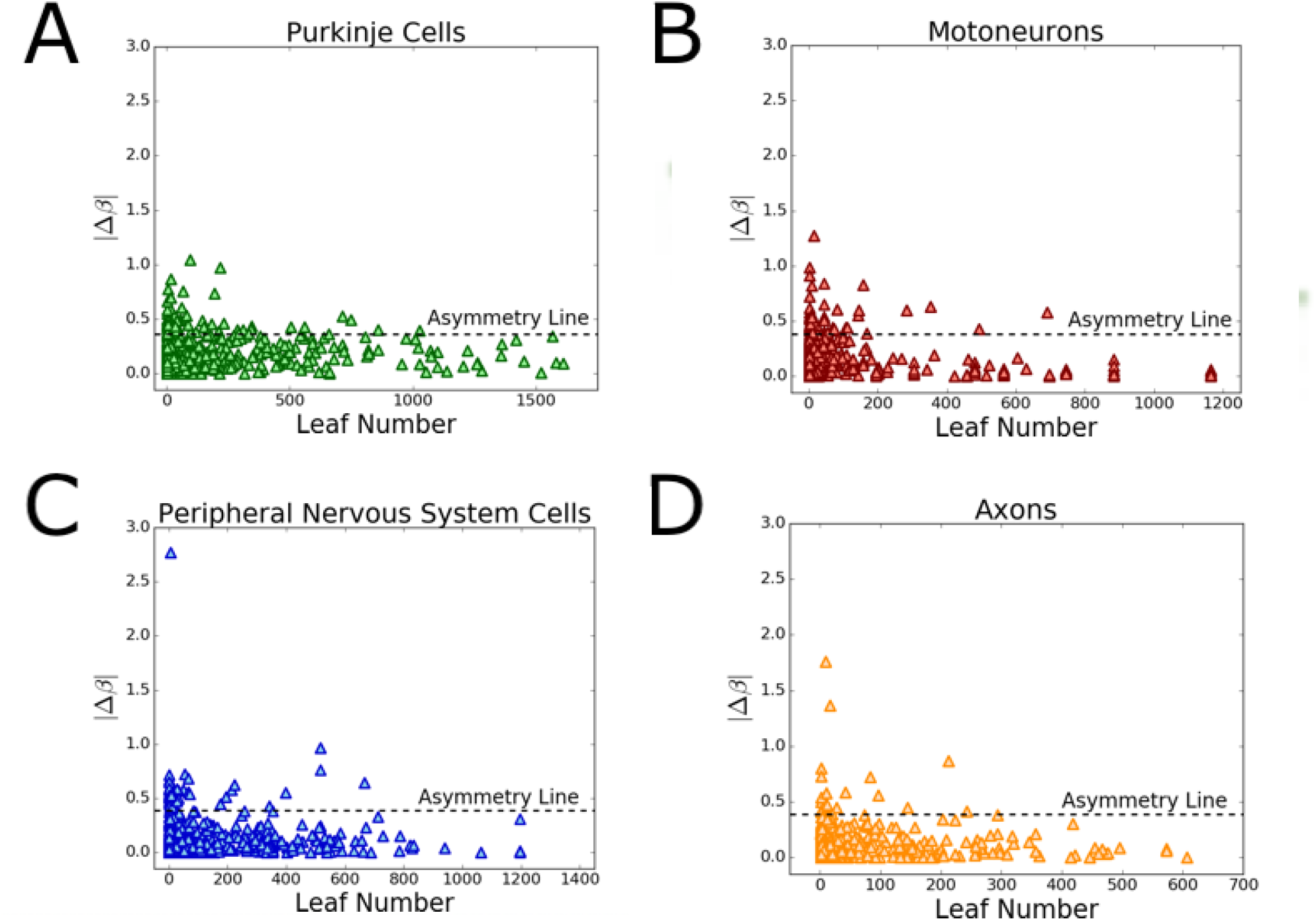
Plots of Degree of Asymmetry vs Leaf Number for A) Purkinje Cells, (B) Motoneurons, (C) Peripheral Nervous System Cells, and (D) Axons

Note that there is an analogous formulation for the difference scale factor for length, 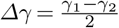. In our analysis, we fix the length scale factor, 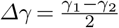, to always be positive. This enforces the following sign convention on the difference scale factor for radius. Consequently, when *Δβ >* 0, one child branch will be both wider and longer than the other child branch. When *Δβ <* 0, one child branch will be wider and shorter than the other child branch. These two scenarios correspond to *positive* and *negative* asymmetric branching and provide a visual way to interpret our results. Here, we focus on branch width rather than length, meaning our results are meaningful in terms of the magnitude but not the direction of asymmetry. For the length scaling to be correctly interpreted, we need to use an alternative [18, 30–32] labeling scheme for branching networks, such as Horton-Strahler labeling. We expand upon this in the Discussion section.

## 3 Methods

We use the method of undetermined Lagrange multipliers to optimize cost functions with varying constraints[33]. When we perform this optimization, we arrive at equations that relate the two radius scaling ratios to each other raised to some scaling exponent (as in Equation 2.9) and corresponding to a generalized conservation rule. We minimize the function by differentiating with respect to each scaling ratio and setting the result equal to zero to solve for the multiplier. Using the fact that the multiplier is constant at each generation *k*, we set _*λk*_ = *λ*_*k*+1_to solve for the resultant equation. More details on each of these calculations can be found in the Appendix.

To test the theoretical predictions and model, we compared the results to data from NeuroMorpho.Org - an online database with digital reconstructions from a wide range of species [26]. These reconstructions are obtained by manually tracing neuron image stacks using computational methods, some manual and some automatic, obtained using microscopic and staining techniques for in vitro neurons and slicing at regular intervals. This database provides 3D reconstruction data that are organized in text files that specify a pixel ID label for each point, the x,y,z spatial coordinates, the radius of the fiber at each point, and a parent pixel ID that refers to the adjacent pixel previously labelled. The scaling ratios for radius and length can be obtained by organizing this data in terms of branches. This is accomplished by finding the pixels at which the difference between the child pixel ID and the parent pixel ID is greater than 2, which can be defined as branching points. Based on the branching points, a branch ID and parent branch ID can be assigned to each of the pixels. The radius can be extracted from each of the branches by taking each of the radius values in each branch and averaging them. The length of each branch can be extracted by summing up the Euclidean distances between each of the points in the branch. Once the radius and length of each of the branches is found, the scaling ratios are computed by dividing the daughter radius by the corresponding value for the parent branch. We can identify the branches that have the same parent to find the two daughters. To extract the scaling exponent P as defined in Equation 2.9, we use the fsolve function in the python library SciPy to numerically solve for the roots of the equation 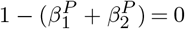.

We look at neuron reconstructions from both axons and dendrites for diverse cell types, brain regions, and species. Due to the small size of axons and the limited resolution of images, the data available on NeuroMorpho.Org are limited in scope. The axon reconstruction data were taken from the following species: fruit flies [34], dragonflies [35], crabs [36], chickens [37], and rats [38]. The neurons were taken from a range of brain regions: the midbrain, the hippocampus, the antennal lobe, the optic lobe, and the ventral nerve cord.

The Purkinje cells are from mice [39–42], rats [42–44], and guinea pigs *Cavia porcellus* [45]. The motoneurons are from zebrafish [46, 47], turtles [48], mice [49], rats [50], rabbits [51], and cats [52].

To study peripheral nervous system (PNS) neurons, we sampled from reconstruction data that was labelled by region on NeuroMorpho.Org. This data was taken from fruit flies [53–55] and mice [56–59] and includes dendritic arborizations, sensory neurons, somatic neurons, and touch receptors.

The scaling ratio data were filtered to remove all daughter pairs where the scaling ratio corresponding to either daughter is equal to 1.0; these values likely occur due to the resolution limit of the image where the radius of both the daughter and the parent branches are equal to the pixel size. Since these values contribute artifacts to the distributions extracted from the data, we remove them from the final dataset.

## 4 Results

Here, we present our results and compare our theoretical predictions with empirical data.

### (a) Theory Results

From the general cost function as described in Equation 2.1, we derive a suite of predictions for scaling relationships. Through this suite of mathematical relationships, we can use optimization to derive powers and corresponding scaling ratios associated with each neuronal function and mechanism. Table 1 summarizes the results of these optimizations. More details on the calculations are in the Appendix.

**Table 1.** Theoretical Predictions for scaling exponents for different functions and comparisons to the median values in the data. The first column is the function that is minimized, either *P* or *T*, as obtained by varying *α* in Equation 2.1. The second column represents a constraint function or a quantity that is held fixed in the optimization. The third column is the result of the minimization using the method of undetermined Lagrange multipliers, and the fourth column is the scaling exponent inferred from these results, which we can compare to the median in the data, including a 95% confidence interval shown in the sixth column.

| Minimize | Constraint | Result | Exponent | Corresponding cell type | Data median (95%CI) |
| --- | --- | --- | --- | --- | --- |
| $P$ | $V$ | $\beta_1^2 + \beta_2^2 = 1$ | 2 | Purkinje cell dendrites | 2.14 (2.03-2.27) |
| $T_{avg,unmyel}$ | $V$ | $\beta_1^{5/2} + \beta_2^{5/2} = 1$ | 5/2 | PNS neurons | 2.98 (2.76-3.16) |
| $T_{max,unmyel}$ | $V$ | $\beta_1^{5/2} = 1$ | 5/2 | PNS neurons | 2.98 (2.76-3.16) |
| $T_{min,unmyel}$ | $V$ | $\beta_2^{5/2} = 1$ | 5/2 | PNS neurons | 2.98 (2.76-3.16) |
| $T_{avg,myel}$ | $V$ | $\beta_1^3 + \beta_2^3 = 1$ | 3 | Axons | 2.96 (2.66-3.23) |
| $T_{max,myel}$ | $V$ | $\beta_1^3 = 1$ | 3 | Axons | 2.96 (2.66-3.23) |
| $T_{min,unmyel}$ | $V$ | $\beta_2^3 = 1$ | 3 | Axons | 2.96 (2.66-3.23) |
| $P$ | $T_{tot,unmyel}$ | $\beta_1^{3/4} + \beta_2^{3/4} = 1$ | 3/4 | Asymmetric motoneurons | 0.90 (0.82-1.08) |
| $P$ | $T_{max,unmyel}$ | $\beta_1^{3/2} = 1$ | 3/2 | Motoneurons | 1.39 (1.35-1.42) |
| $P$ | $T_{min,unmyel}$ | $\beta_2^{3/2} = 1$ | 3/2 | Motoneurons | 1.39 (1.35-1.42) |

### (b) Data Results

Here, we compare the theoretical predictions to empirical results, including histograms showing distributions of scaling exponents and the relationship between asymmetry and network level, or distance from the soma. The scaling exponent data was restricted to values above 0. As in the neuroscience literature [29], we compared the median values in the data to the theoretical predictions. The colored dotted lines show the theoretical predictions for each of comparison to the median as well as the relative peaks in the data.

#### i Asymmetry Distributions Across Branching Generations

Figure 3 shows the distributions of asymmetry in the branching junctions of each of the neurite types, where degree of asymmetry is represented as the difference scale factor, *Δβ*. The value of *μ* is the mean of the data, and *σ* is the standard deviation. For all of the neurite types, there is a normal distribution of asymmetry factors, with a peak at the symmetric case, where *Δβ* = 0. Purkinje cells show the least asymmetry, as *σ* is the smallest, which is consistent with expectations based on the visual symmetries in their branching architecture.

Figure 4 shows plots relating the degree of asymmetry to the Leaf Number, where the smaller leaf numbers are the tips closest to the synapses, as illustrated in Figure 3. Here, we focus on the magnitude of the difference scale rather than the direction of asymmetry, defined by the absolute value of the difference scale factor *Δβ*, as defined previously.

The horizontal dashed line is what we define as a cutoff for the asymmetry line— a difference scale factor *Δβ* more than two standard deviations away from the mean (at which symmetry occurs)— that shows the division between the symmetric and asymmetric modes. We observe that the most asymmetric branching junctions, or those that occur above the asymmetry line, occur at lower leaf numbers, or closer to the synapses in the neuronal network.

#### ii Overall Network Power Distributions

Figure 5 shows the distributions of scaling exponents solved from the data. These show general network-wide trends in branching, and the corresponding solid black lines are the medians in the data. The medians in the Purkinje cell and motoneuron scaling exponent data correspond to the theoretical predictions for the functions minimizing power and the medians in the axons and PNS neuron scaling exponent data correspond to the theoretical predictions for the functions minimizing conduction time delay. The motoneuron data correspond to the prediction for the function that includes conduction time delay as a biophysical constraint, while the Purkinje cell data correspond to the prediction for the function that includes a material constraint.

**Fig 5.**
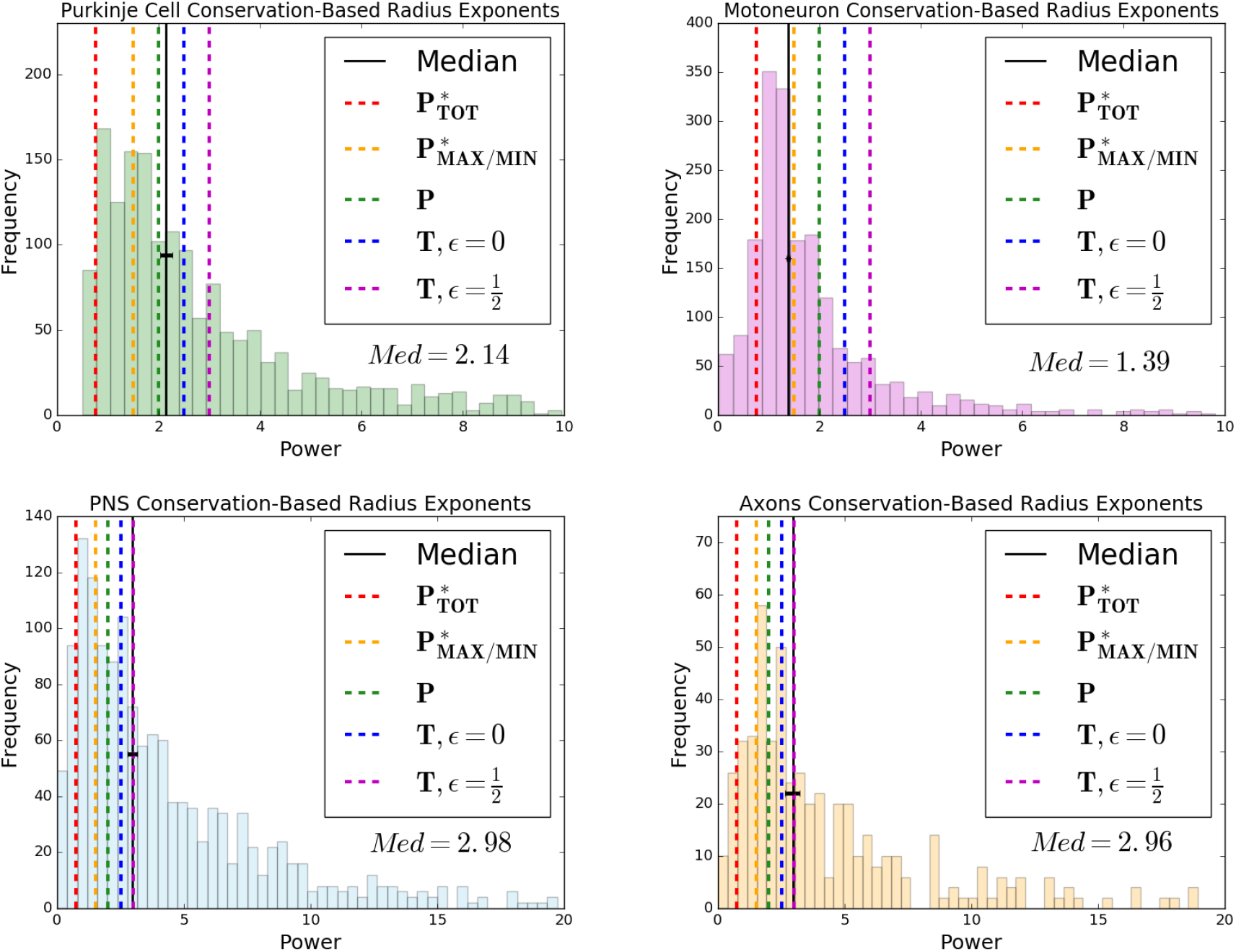
Branching Scaling Exponent Data

#### iii Symmetric Versus Asymmetric Motoneuron Branching Junctions

Although the general network-wide trends are useful, we also split the data based on degree of asymmetry. We split the motoneuron data into symmetric and asymmetric branches, where the difference scale factor for the symmetric data fall within two standard deviations from 0. Analyzing the data separately in Figure 6, we find different median powers that correspond to theoretical predictions from different functions. The scaling exponent data for asymmetric branching junctions in motoneurons corresponds to the theoretical prediction for the function that interprets the conduction time delay as a sum of all possible paths, while the data for the symmetric branching junctions correspond to the prediction for the function that considers one optimal path.

**Fig 6.**
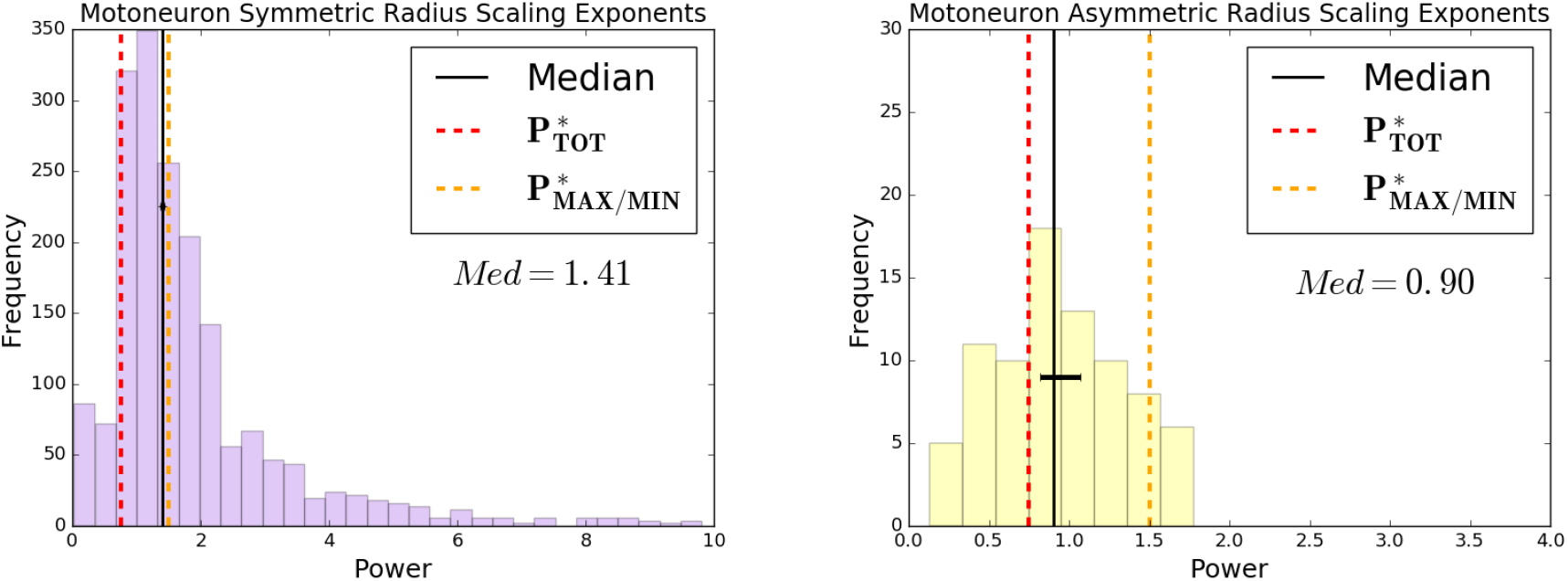
Symmetric and Asymmetric Scaling Exponent Data for Motoneuron Branching Junctions

## 5 Discussion

Asymmetric branching in neurons gives rise to multiple possible paths from the soma to the synapses and vice versa. Although a symmetric branching network model can provide insights into the connections between branching patterns and functional principles of neurons, it obscures key features of these networks, such as large differences in path lengths from soma to tips and how those contribute to the functionality of neurons. The introduction of asymmetric branching to our model gives rise to multiple possible interpretations of the conduction time delay term, one that focuses on optimizing the path associated with either the maximum or minimum conduction time delay in the network, and another that takes into consideration the sum of all paths in the network. Notably, the mathematical results of the optimization of the models are the same for the symmetric model [18] and the maximum or minimum interpretation of conduction time delay in the asymmetric case. However, the total path interpretation of the conduction time delay leads to different results for the function minimizing power with a time delay constraint. Moreover, for motoneurons, we find that splitting the scaling exponent data into the most symmetric and the most asymmetric data leads to different median values that correspond to theoretical predictions for different interpretations of this constraint, which also correspond to different regions in the cell relative to the soma and synapses. The median for asymmetric junctions corresponds to the theoretical prediction with the total path length interpretation of the conduction time delay constraint, suggesting that the whole network— rather than just one optimal path— is important for asymmetric branching junctions. The symmetric model obscures this distinction, and thus our comparisons of asymmetric and symmetric branching junctions lead us to look more closely at the position of branching junctions relative to the soma or the synapses, and whether there is any connection between this position and asymmetry.

We define this position using the key measure of leaf number. The distributions of leaf numbers shown in Figure 4 reflect what we know about differences in the structure of axons and dendrites. The maximum leaf number for axons is around 600, while it is around 1200-1600 for dendrites, thus reflecting the longer single parent branch for axons versus the more extensive branching and the greater number of branching generations for dendrites. Moreover, there are significant differences observed among different types of dendritic structures. The maximum leaf numbers for motoneurons and PNS neurons are around 1200, while it is around 1600 for Purkinje cells. We hypothesize that these differences might be due to differences in extracellular environments; motoneurons and PNS neurons are both part of the sensorimotor circuits that are localized in distal parts of the body, while Purkinje cells are located in the cerebellum within the brain itself. As dendrite branch formation is controlled by guidance cues in the environment that trigger complex intracellular signalling cascades and lead to protrusions [60], the vastly disparate biological environments and extrinsic cues likely greatly influence the extent of dendritic branching in these different cell types.

Although our analysis of the correspondences across cell types focuses on median values in the data, as seen in other biological networks [23], we find much more variance at the local level. In neuroscience, theoretical and experimental work has shown that motoneurons grow in a roughly self-referential manner and their basic structure and branching points are predetermined. However, environmental cues and activity-dependent behavior cause local changes in morphology [60, 61]. Here, we are able to observe that the median scaling exponents differ significantly for symmetric and asymmetric branching in motoneurons, corresponding to different theoretical predictions. It is possible that the variance in symmetry of branching junctions might account for the wide distribution of scaling exponents for each cell type. The distributions of scaling exponents have a wide variance and multiple local peaks. Although we map the median scaling exponent to the closest theoretical prediction in this analysis, it is possible that there are multiple functional principles at play at different localized regions of the cells, corresponding to the peaks observed across the distribution. Moreover, we observe a correspondence between asymmetry of branching junctions and their relative position in a neurite, whether they are closer to the tips or to the soma. At the tips, where the leaf number is closest to zero, the branching junctions can be either extremely symmetric or extremely asymmetric.

Importantly, the most asymmetric branching junctions always occur at the tips. In contrast, the branching junctions that occur closest to the soma all fall under the symmetric type, where the *Δ β* value is within two standard deviations from the symmetric case. Thus, we observe **two different symmetry/asymmetry regimes, with a shift from the most symmetric branching at the soma to an increased number of asymmetric branching junctions at the tips**.

Because the tips of axons and dendrites are closest to the synapses, this suggests that the asymmetry might have to do with the forming of actual connections at the tips. This is consistent with existing knowledge that the branching of axons and dendrites is determined by synapses; new branches are formed preferentially near the synapses [11]. Moreover, previous studies have shown that there are activity-dependent changes in morphology of motoneurons [62]. Studies of other biological networks have also shown that they are robust to damage and changes in the environment, developing corresponding changes in morphology to adapt to environments [63]. It is possible that the difference in power observed at the tip is due to changes as a result of activity-dependent behavior such as synaptic formation and pruning. This is consistent with empirically-informed mathematical models that describe the elongation of neurites as an extension of the cytoskeleton, where the most active building blocks (microtubules) are located at the distal portions and tips, making them more susceptible to developmental variation based on environmental cues [64]. Moreover, the fact that for asymmetric branching junctions, the interpretation of conduction time delay as a sum of all possible paths rather than one optimal path supports the notion that asymmetric branching is connected to plasticity in the network. These asymmetric branching patterns are determined by the sum of paths across the whole network, suggesting that the whole network is optimized in a way that is robust to damage in single paths and such that the whole network is optimized to make as many synaptic connections with other neurites as possible.

In this analysis, we have chosen to focus on radius scaling ratios and asymmetries that occur in the width of daughter branches. Although length asymmetry might provide additional insights into the properties of these networks, the branch length measurements are not accurately characterized, as also previously reported for vascular scaling [30] as well as other types of plant and animal networks [31, 32]. Recent work suggests Horton-Strahler labeling — where the first level begins at the tips, and higher levels are determined when two branches of the same level combine — may yield better estimates of branch length scaling, as it has been previously applied to neurons and other biological networks [65–67]. In future work, we plan to investigate how this alternative labeling scheme for branch lengths compares with theoretical predictions derived using our framework. If we are able to obtain meaningful results from the analysis of length scaling ratios, the direction of asymmetry and the distinction between the two types—*positive* and *negative* asymmetry— will be an important consideration in addition to the magnitude which we focused on here.

Moreover, we aim to formulate a new constraint that relates to the way in which neurons fill space. So far, our optimization considers only intrinsic properties of neurons without explicitly accounting for: 1. interactions amongst neurons, 2. electrical activity that might strengthen or prune synapses, and 3. environmental chemoattractants and chemorepellants that might shape the growth and development of neurons, particularly in relation to their length. Adding this interaction term might lead us to understand length scaling ratios more.

Future studies have the potential to illuminate the function of asymmetry in neuron plasticity by analyzing in-vivo neuron image data taken across stages of development. Long term, a greater understanding of the details of the asymmetries observed within and among neurites and single cells may help pave the way to understanding lateral asymmetries in the brain and the structure-function correspondence.

In conclusion, we find that our asymmetric branching model for axons and dendrites brings to light the importance of considering all possible paths from the synapse to the soma rather than one optimal path. While this distinction does not affect the predictions for functions that minimize conduction time delay, they alter the predictions for the functions that minimize power and fix conduction time delay as a constraint. For motoneurons, the different interpretations of conduction time delay correspond to the median in the scaling exponent data of different types of branching junctions. The symmetric branching junctions agree with the predictions focusing on one optimal path, while the asymmetric branching junctions agree with the predictions that take all paths into consideration.

Moreover, the asymmetric branching junctions are localized closer to the synapses, suggesting that there is some connection between asymmetric branching and environmental factors, plasticity, and whole network robustness. This distinction between predictions for asymmetric and symmetric branching is observed only when time delay is a constraint (as opposed to a function to be minimized) and for motoneuron dendrites (but not axons). This is consistent with the notion that dendrites, in contrast to axons, are shorter with more extensive branching that allows them to connect to multiple other neurons [10]. Our results support the notion that the whole network with its various paths—rather than simply optimal paths—are important factors governing the structures of these dendrites. Dendrite branches must reach multiple potential synaptic targets, and these synaptic connections are constantly evolving, forming, and pruning. This asymmetric branching framework is necessary in order to study and reason about these features of the network.

## Data Accessibility

Data are available at www.neuromorpho.org. The code used to analyze this data as well as details about the file names of the cells we analyzed, including the cell type, region, species, and archive from which the data were taken can be accessed at the GitHub repository https://github.com/pahelidc/scalingtheoryneurons

## Authors’ Contributions

P.D.C. contributed to conceptualization, writing the code, funding acquisition, formal analysis, writing the original draft, drawing (Figures 1 and 2), and reviewing and editing. A.B.B contributed to conceptualization and reviewing and editing.

S. M. contributed to writing the code and generating the plots in Figures 3 and 4. V.M.S contributed to conceptualization, reviewing and editing, and supervision.

