## Appendix for "Neuronal Branching is Increasingly Asymmetric Near Synapses, Potentially Enabling Plasticity While Minimizing Energy Dissipation and Conduction Time"

### A Appendix 1

In Table 1 in the main text, we show a series of results calculated from specific cases of the general cost function in Equation 2.1. Here, we will show the calculations of these results in more detail.

#### (a) Power Minimization with Fixed Volume

The simplest case for this type of calculation is the optimization of a cost function that minimizes power lost due to dissipation ($\alpha = 1$ in Equation 2.1) with a material cost constraint. We have represented the material cost constraint here as the total network volume, and this is a quantity that we hold to be fixed in the optimization.

The specific function we are minimizing here is:

$$\begin{aligned} P_{TOT} = I_0^2 R_{N,TOT} \sum_{k=0}^{N} \left( \prod_{j=k}^{N-1} \left[ \frac{\beta_{j,1}^2}{\gamma_{j,1}} + \frac{\beta_{j,2}^2}{\gamma_{j,2}} \right] \right) + \lambda V_{N,TOT} \sum_{k=0}^{N} \left( \prod_{j=k}^{N-1} \left[ \beta_{j,1}^2 \gamma_{j,1} + \beta_{j,2}^2 \gamma_{j,2} \right]^{-1} \right) \\ + \lambda_m m_c + \sum_{k=0}^{N} \left( \lambda_k l_{N,TOT}^d \prod_{j=k}^{N-1} \left[ \gamma_{j,1}^d + \gamma_{j,2}^d \right]^{-1} \right) \end{aligned} \tag{A 1}$$

We begin the optimization by taking the derivative of the function with respect to $\beta_{i,1}$ and $\beta_{i,2}$ and setting them equal to 0.

$$\frac{\partial P_{TOT}}{\partial \beta_{i,1}} = \sum_{k=0}^{N} \left( \frac{2 I_0^2 R_{N,TOT} \beta_{i,1}}{\gamma_{i,1}} \prod_{j=k, j\neq i}^{N-1} \left[ \frac{\beta_{j,1}^2}{\gamma_{j,1}} + \frac{\beta_{j,2}^2}{\gamma_{j,2}} \right] - \frac{2\lambda V_{N,TOT} \beta_{i,1} \gamma_{i,1}}{\beta_{i,1}^2 \gamma_{i,1} + \beta_{i,2}^2 \gamma_{i,2}} \prod_{j=k, j\neq i}^{N-1} \left[ \beta_{j,1}^2 \gamma_{j,1} + \beta_{j,2}^2 \gamma_{j,2} \right]^{-1} \right) \tag{A 2}$$

$$\frac{\partial P_{TOT}}{\partial \beta_{i,2}} = \sum_{k=0}^{N} \left( \frac{2 I_0^2 R_{N,TOT} \beta_{i,2}}{\gamma_{i,2}} \prod_{j=k, j\neq i}^{N-1} \left[ \frac{\beta_{j,1}^2}{\gamma_{j,1}} + \frac{\beta_{j,2}^2}{\gamma_{j,2}} \right] - \frac{2\lambda V_{N,TOT} \beta_{i,2} \gamma_{i,2}}{\beta_{i,1}^2 \gamma_{i,1} + \beta_{i,2}^2 \gamma_{i,2}} \prod_{j=k, j\neq i}^{N-1} \left[ \beta_{j,1}^2 \gamma_{j,1} + \beta_{j,2}^2 \gamma_{j,2} \right]^{-1} \right) \tag{A 3}$$

Since both of these equations, A.2 and A.3, are equal to 0, we can simplify this expression by adding a linear combination of these equations, which is also equal to 0. We will multiply A.2 by $\beta_{i,1}$ and multiply A.3 by $\beta_{i,2}$, arriving at the following expression.

$$0 = \sum_{k=0}^{N} \left( 2 I_0^2 R_{N,TOT} \prod_{j=k}^{N-1} \left[ \frac{\beta_{j,1}^2}{\gamma_{j,1}} + \frac{\beta_{j,2}^2}{\gamma_{j,2}} \right] - 2\lambda V_{N,TOT} \prod_{j=k}^{N-1} \left[ \beta_{j,1}^2 \gamma_{j,1} + \beta_{j,2}^2 \gamma_{j,2} \right]^{-1} \right) \tag{A 4}$$

Since all of these terms are positive biological quantities and because the sum of the terms is equal to 0, then each of the individual terms in the sum must be equal to zero. Using this fact, we can solve for an expression for $\lambda$.

$$\lambda = \frac{I_0^2 R_{N,TOT}}{V_{N,TOT}} \prod_{j=k}^{N-1} \left[ \beta_{j,1}^2 \gamma_{j,1} + \beta_{j,2}^2 \gamma_{j,2} \right] \left[ \frac{\beta_{j,1}^2}{\gamma_{j,1}} + \frac{\beta_{j,2}^2}{\gamma_{j,2}} \right] \tag{A 5}$$

If we plug this expression for $\lambda$ back into A.2, we can simplify the expression to the following:

$$\gamma_{i,1} = \gamma_{i,2} \tag{A 6}$$

Now, using the fact that $\lambda$ is a constant and thus stays the same across generations; that is, $\lambda_k = \lambda_{k+1}$, we arrive at the following expression:

$$\left[ \beta_{k,1}^2 \gamma_{k,1} + \beta_{k,2}^2 \gamma_{k,2} \right] \left[ \frac{\beta_{k,1}^2}{\gamma_{k,1}} + \frac{\beta_{k,2}^2}{\gamma_{k,2}} \right] = 1 \tag{A 7}$$

Substituting A.6 in A.7 and simplifying, we arrive at

$$\beta_{k,1}^2 + \beta_{k,2}^2 = 1 \tag{A 8}$$

That is, the two scaling ratios are raised to a scaling exponent of 2. Note that in this case, we can rewrite the equation by expanding $\beta_{k,1}$ and $\beta_{k,2}$ based on their definitions, $\beta_{k,1} = \frac{r_{k+1,1}}{r_k}$ and $\beta_{k,2} = \frac{r_{k+1,2}}{r_k}$.

$$\left(\frac{r_{k+1,1}}{r_k}\right)^2 + \left(\frac{r_{k+1,2}}{r_k}\right)^2 = 1 \tag{A 9}$$

This leads to the following relationship between the radii:

$$r_{k+1,1}^2 + r_{k+1,2}^2 = r_k^2 \tag{A 10}$$

Note that this implies that the sum of the cross-sectional areas of the two daughter branches is equal to the cross-sectional area of the parent branch (scaling this equation by $\pi$ would give this relationship). This implies that our scaling exponent relationship is **area-preserving** in this case. There is an analogous area-preserving case observed in studies of cardiovascular branching blood vessels[1].

#### (b) Time Minimization, Umyelinated

The calculations involving optimization of conduction time delay are more complicated because in an asymmetric branching network, there are multiple possible paths from the soma to the synapses. Thus, different interpretations correspond to different optimization problems.

##### i Average Path Interpretation

For the average interpretation, the idea is that for a bifurcating branching network, there is an average time at each level, divided by the total number of branches, $2^k$. (This can be generalized for other types of networks with branching ratio $n$ as $n^k$). Thus, we can define the function we are optimizing as follows:

$$\begin{aligned} T = T_{TOT} \sum_{k=0}^{N} \frac{1}{2^k} \left( \prod_{j=k}^{N-1} \left[ \frac{\gamma_{j,1}}{\beta_{j,1}^{1/2}} + \frac{\gamma_{j,2}}{\beta_{j,2}^{1/2}} \right] \right) + \lambda V_{N,TOT} \sum_{k=0}^{N} \left( \prod_{j=k}^{N-1} \left[ \beta_{j,1}^2 \gamma_{j,1} + \beta_{j,2}^2 \gamma_{j,2} \right]^{-1} \right) \\ + \lambda_m m_c + \sum_{k=0}^{N} \left( \lambda_k l_{N,TOT}^d \prod_{j=k}^{N-1} \left[ \gamma_{j,1}^d + \gamma_{j,2}^d \right]^{-1} \right) \end{aligned} \tag{A 11}$$

Following the same steps for the optimization as in section a, we arrive at

$$\beta_{k,1}^{5/2} + \beta_{k,2}^{5/2} = 1 \tag{A 12}$$

That is, the two scaling ratios are raised to a scaling exponent of 5/2. As we saw before, we can rewrite this relationship as follows.

$$r_{k+1,1}^{5/2} + r_{k+1,2}^{5/2} = r_k^{5/2} \tag{A 13}$$

While the previous case was area-preserving, this case is **area-increasing**.

##### ii Maximum/Minimum Path Interpretation

For both the maximum and minimum path interpretations of conduction time delay, we focus on just one of the $\beta$ values, the larger one, $\beta_1$ corresponding to the minimum time delay path (as the velocity is the greatest) and the smaller one, $\beta_2$ for maximum time delay path (as the velocity is the smallest). For simplicity, we will focus on the calculation minimizing the conduction time delay of the optimal path using $\beta_1$, as the calculations are mathematically equivalent.

Thus, we can define the function we are optimizing as follows:

$$\begin{aligned} T = T_{TOT} \sum_{k=0}^{N} \left( \prod_{j=k}^{N-1} \frac{\gamma_{j,1}}{\beta_{j,1}^{1/2}} \right) + \lambda V_{N,TOT} \sum_{k=0}^{N} \left( \prod_{j=k}^{N-1} \left[ \beta_{j,1}^2 \gamma_{j,1} + \beta_{j,2}^2 \gamma_{j,2} \right]^{-1} \right) \\ + \lambda_m m_c + \sum_{k=0}^{N} \left( \lambda_k l_{N,TOT}^d \prod_{j=k}^{N-1} \left[ \gamma_{j,1}^d + \gamma_{j,2}^d \right]^{-1} \right) \end{aligned} \tag{A 14}$$

Note that here, since we are focusing on $\beta_1$, corresponding to the maximum velocity or the minimum conduction time delay, we are minimizing the minimum path here. We take the derivative of the function with respect to $\beta_{i,1}$ and $\beta_{i,2}$ and set them equal to 0.

$$\frac{\partial T}{\partial \beta_{i,1}} = \sum_{k=0}^{N} \left( \frac{T_{TOT}}{2} \frac{\beta_{i,1}^{-1/2}}{\gamma_{i,1}} \prod_{j=k, j\neq i}^{N-1} \frac{\beta_{j,1}^{1/2}}{\gamma_{j,1}} - \frac{2\lambda V_{N,TOT}\beta_{i,1}\gamma_{i,1}}{\beta_{i,1}^2\gamma_{i,1} + \beta_{i,2}^2\gamma_{i,2}} \prod_{j=k, j\neq i}^{N-1} \left[\beta_{j,1}^2\gamma_{j,1} + \beta_{j,2}^2\gamma_{j,2}\right]^{-1} \right) \tag{A 15}$$

$$\frac{\partial T}{\partial \beta_{i,1}} = -\frac{2\lambda V_{N,TOT}\beta_{i,2}\gamma_{i,2}}{\beta_{i,1}^2\gamma_{i,1} + \beta_{i,2}^2\gamma_{i,2}} \prod_{j=k, j\neq i}^{N-1} \left[\beta_{j,1}^2\gamma_{j,1} + \beta_{j,2}^2\gamma_{j,2}\right]^{-1} \Bigg) \tag{A 16}$$

Note that since $\beta_{i,2}$ does not appear in the conduction time delay term being minimized, the first term disappears when taking the derivative with respect to $\beta_{i,2}$. As before, since both of these equations, A.15 and A.16, are equal to 0, we can simplify this expression by adding a linear combination of these equations, which is also equal to 0. We will multiply A.15 by $\beta_{i,1}$ and multiply A.16 by $\beta_{i,2}$, arriving at the following expression.

$$0 = \sum_{k=0}^{N} \left( \frac{T_{TOT}}{2} \prod_{j=k}^{N-1} \frac{\beta_{j,1}^{1/2}}{\gamma_{j,1}} - 2\lambda V_{N,TOT} \prod_{j=k}^{N-1} \left[\beta_{j,1}^2\gamma_{j,1} + \beta_{j,2}^2\gamma_{j,2}\right]^{-1} \right) \tag{A 17}$$

As before, we can solve for an expression for $\lambda$.

$$\lambda = \frac{T_{TOT}}{4V_{N,TOT}} \prod_{j=k}^{N-1} \frac{\beta_{j,1}^{1/2}}{\gamma_{j,1}} \left[\frac{\beta_{j,1}^2}{\gamma_{j,1}} + \frac{\beta_{j,2}^2}{\gamma_{j,2}}\right] \tag{A 18}$$

If we plug this expression for $\lambda$ back into A.15, we can simplify the expression to the following:

$$\beta_{i,2}^2\gamma_{i,2} = 0 \tag{A 19}$$

Now, using the fact that $\lambda$ is a constant and thus stays the same across generations; that is, $\lambda_k = \lambda_{k+1}$, we arrive at the following expression:

$$\frac{\beta_{k,1}^{1/2}}{\gamma_{k,1}} \left[\beta_{k,1}^2\gamma_{k,1} + \beta_{k,2}^2\gamma_{k,2}\right] = 1 \tag{A 20}$$

By substituting A.19 in A.20 and simplifying, we arrive at

$$\beta_{k,1}^{5/2} = 1 \tag{A 21}$$

Here, we find that the scaling exponent, which only applies to $\beta_1$, is 5/2. Repeating this calculation with $\beta_2$, focusing on the maximum path, would yield the equation $\beta_{k,2}^{5/2} = 1$. As above, substituting the expressions for the scaling ratios, we can rewrite this as

$$r_{k+1,1}^{5/2} = r_k^{5/2} \tag{A 22}$$

This is for the minimum path calculation. Analogously, if we focus on the maximum path, we can write this equation as $r_{k+1,1}^{5/2} = r_k^{5/2}$. For either case, focusing on the relationship between the parent and the selected daughter branch, we see that the relationship is **area-increasing**.

#### (c) Time Minimization, Myelinated

Here, we repeat the above calculations for the myelinated case, where the velocity is proportional to the radius of each branch rather than the square root of the radius. Thus, we begin with a similar function but with different powers for $\beta$ in the conduction time delay term.

##### i Average Path Interpretation

As above, for the average interpretation, by the total number of branches, $2^k$. Thus, we can define the function we are optimizing as follows:

$$T = T_{TOT}\sum_{k=0}^{N}\frac{1}{2^k}\left(\prod_{j=k}^{N-1}\left[\frac{\gamma_{j,1}}{\beta_{j,1}}+\frac{\gamma_{j,2}}{\beta_{j,2}}\right]\right)+\lambda V_{N,TOT}\sum_{k=0}^{N}\left(\prod_{j=k}^{N-1}\left[\beta_{j,1}^2\gamma_{j,1}+\beta_{j,2}^2\gamma_{j,2}\right]^{-1}\right)$$
$$+\lambda_m m_c+\sum_{k=0}^{N}\left(\lambda_k l_{N,TOT}^d\prod_{j=k}^{N-1}\left[\gamma_{j,1}^d+\gamma_{j,2}^d\right]^{-1}\right) \quad \text{(A 23)}$$

Following the same steps for the optimization as in sections a and b, we arrive at

$$\beta_{k,1}^3+\beta_{k,2}^3=1 \quad \text{(A 24)}$$

That is, the two scaling ratios are raised to a scaling exponent of 3, and this relationship is **area-increasing**. Note that this is the same as the area-increasing relationship pretended for branching blood vessels [1].

##### ii Maximum/Minimum Path Interpretation

As before, we optimize the function focusing on the path calculated with the larger scaling ratio, $\beta_1$, corresponding to the path of minimum conduction time delay.

The function we are optimizing for the myelinated case is as follows:

$$T = T_{TOT}\sum_{k=0}^{N}\left(\prod_{j=k}^{N-1}\frac{\gamma_{j,1}}{\beta_{j,1}}\right)+\lambda V_{N,TOT}\sum_{k=0}^{N}\left(\prod_{j=k}^{N-1}\left[\beta_{j,1}^2\gamma_{j,1}+\beta_{j,2}^2\gamma_{j,2}\right]^{-1}\right)$$
$$+\lambda_m m_c+\sum_{k=0}^{N}\left(\lambda_k l_{N,TOT}^d\prod_{j=k}^{N-1}\left[\gamma_{j,1}^d+\gamma_{j,2}^d\right]^{-1}\right) \quad \text{(A 25)}$$

Following the same steps for the optimization as in sections a and b, we arrive at

$$\beta_{k,1}^3=1 \quad \text{(A 26)}$$

Here, we find that the scaling exponent, which only applies to $\beta_1$, is 3, for the minimum path. Repeating this calculation with $\beta_2$ would yield the equation $\beta_{k,2}^3=1$ for the maximum path. As before, this relationship is **area-increasing**.

#### (d) Power Minimization with Fixed Time Delay

Here, we repeat these calculations for the case where the constraint is conduction time delay rather than volume.

##### i Total Paths Interpretation

For the total paths interpretation of the conduction time delay, we are optimizing the following function:

$$P^* = I_0^2 R_{N,TOT}\sum_{k=0}^{N}\left(\prod_{j=k}^{N-1}\left[\frac{\beta_{j,1}^2}{\gamma_{j,1}}+\frac{\beta_{j,2}^2}{\gamma_{j,2}}\right]\right)-\lambda T_{TOT}\sum_{k=0}^{N}\left(\prod_{j=k}^{N-1}\left[\frac{\gamma_{j,1}}{\beta_{j,1}^{1/2}}+\frac{\gamma_{j,2}}{\beta_{j,2}^{1/2}}\right]\right)$$
$$+\lambda_m m_c+\sum_{k=0}^{N}\left(\lambda_k l_{N,TOT}^d\prod_{j=k}^{N-1}\left[\gamma_{j,1}^d+\gamma_{j,2}^d\right]^{-1}\right) \quad \text{(A 27)}$$

Note that the sign of the constraint function is arbitrary, so we use the negative one to simplify the calculation. We begin the optimization by taking the derivative of the function with respect to $\beta_{i,1}$ and $\beta_{i,2}$ and setting them equal to 0.

$$\frac{\partial P^*}{\partial\beta_{i,1}}=\sum_{k=0}^{N}\left(\frac{2I_0^2R_{N,TOT}\beta_{i,1}}{\gamma_{i,1}}\prod_{j=k,j\neq i}^{N-1}\left[\frac{\beta_{j,1}^2}{\gamma_{j,1}}+\frac{\beta_{j,2}^2}{\gamma_{j,2}}\right]-\frac{T_{TOT}\gamma_{i,1}}{2\beta_{i,1}^{3/2}\left[\frac{\gamma_{j,1}}{\beta_{j,1}}+\frac{\gamma_{j,2}}{\beta_{j,2}}\right]^2}\prod_{j=k,j\neq i}^{N-1}\left[\frac{\gamma_{j,1}}{\beta_{j,1}^{1/2}}+\frac{\gamma_{j,2}}{\beta_{j,2}^{1/2}}\right]^{-1}\right) \quad \text{(A 28)}$$

$$\frac{\partial P^*}{\partial \beta_{i,2}} = \sum_{k=0}^{N}\left(\frac{2I_0^2 R_{N,TOT}\beta_{i,2}}{\gamma_{i,2}}\prod_{j=k,j\neq i}^{N-1}\left[\frac{\beta_{j,1}^2}{\gamma_{j,1}}+\frac{\beta_{j,2}^2}{\gamma_{j,2}}\right] - \frac{T_{TOT}\gamma_{i,2}}{2\beta_{i,2}^{3/2}\left[\frac{\gamma_{j,1}}{\beta_{j,1}}+\frac{\gamma_{j,2}}{\beta_{j,2}}\right]^2}\prod_{j=k,j\neq i}^{N-1}\left[\frac{\gamma_{j,1}}{\beta_{j,1}^{1/2}}+\frac{\gamma_{j,2}}{\beta_{j,2}^{1/2}}\right]^{-1}\right) \tag{A 29}$$

Since both of these equations, A.28 and A.29, are equal to 0, we can simplify this expression by adding a linear combination of these equations, which is also equal to 0. We will multiply A.28 by $\beta_{i,1}$ and multiply A.29 by $\beta_{i,2}$, arriving at the following expression.

$$0 = \sum_{k=0}^{N}\left(2I_0^2 R_{N,TOT}\prod_{j=k}^{N-1}\left[\frac{\beta_{j,1}^2}{\gamma_{j,1}}+\frac{\beta_{j,2}^2}{\gamma_{j,2}}\right] - \lambda\frac{T_{TOT}}{2}\prod_{j=k,j\neq i}^{N-1}\left[\frac{\gamma_{j,1}}{\beta_{j,1}^{1/2}}+\frac{\gamma_{j,2}}{\beta_{j,2}^{1/2}}\right]^{-1}\right) \tag{A 30}$$

Since all of these terms are positive biological quantities and because the sum of the terms is equal to 0, then each of the individual terms in the sum must be equal to zero. Using this fact, we can solve for an expression for $\lambda$.

$$\lambda = \frac{-4I_0^2 R_{N,TOT}}{T_{TOT}}\prod_{j=k}^{N-1}\left[\frac{\beta_{j,1}^2}{\gamma_{j,1}}+\frac{\beta_{j,2}^2}{\gamma_{j,2}}\right]\left[\frac{\gamma_{j,1}}{\beta_{j,1}^{1/2}}+\frac{\gamma_{j,2}}{\beta_{j,2}^{1/2}}\right] \tag{A 31}$$

If we plug this expression for $\lambda$ back into A.28, we can simplify the expression to the following:

$$\frac{\gamma_{i,1}}{\gamma_{i,2}} = \frac{\beta_{i,1}^{5/4}}{\beta_{i,2}^{5/4}} \tag{A 32}$$

Now, using the fact that $\lambda$ is a constant and thus stays the same across generations; that is, $\lambda_k = \lambda_{k+1}$, we arrive at the following expression:

$$\beta_{k,1}^{3/2} + \frac{\beta_{k,1}^2}{\beta_{k,2}^{1/2}}\left[\frac{\gamma_{k,2}}{\gamma_{k,1}}\right] + \frac{\beta_{k,2}^2}{\beta_{k,1}^{1/2}}\left[\frac{\gamma_{i,1}}{\gamma_{i,2}}\right] + \beta_{k,2}^{3/2} = 1 \tag{A 33}$$

Substituting A.32 in A.33, simplifying, and factoring, we arrive at

$$\beta_{k,1}^{3/4} + \beta_{k,2}^{3/4} = 1 \tag{A 34}$$

That is, the two scaling ratios are raised to a scaling exponent of 3/4. As before, we can also represent this expression as

$$r_{k+1,1}^{3/4} + r_{k+1,2}^{3/4} = r_k^{3/4} \tag{A 35}$$

Note that this relationship, unlike the others mentioned above, is **area-decreasing**, as this scaling exponent is less than 2 and the sum of the cross-sectional areas of the daughter branches is less than the cross-sectional area of the parent.

##### ii Maximum/Minimum Path Interpretation

For the maximum/minimum interpretation of the conduction time delay constraint, we are optimizing the following function, focusing on $\beta_1$ (minimum path):

$$P^* = I_0^2 R_{N,TOT}\sum_{k=0}^{N}\left(\prod_{j=k}^{N-1}\left[\frac{\beta_{j,1}^2}{\gamma_{j,1}}+\frac{\beta_{j,2}^2}{\gamma_{j,2}}\right]\right) - \lambda T_{TOT}\sum_{k=0}^{N}\left(\prod_{j=k}^{N-1}\frac{\beta_{j,1}^{1/2}}{\gamma_{j,1}}\right) + \lambda_m m_c + \sum_{k=0}^{N}\left(\lambda_k l_{N,TOT}^d\prod_{j=k}^{N-1}\left[\gamma_{j,1}^d+\gamma_{j,2}^d\right]^{-1}\right) \tag{A 36}$$

Following the same steps for the optimization as in sections a, b, and c, we arrive at

$$\beta_{k,1}^{3/2} = 1 \tag{A 37}$$

Here, we find that the scaling exponent, which only applies to $\beta_1$, is 3/2. Repeating this calculation with $\beta_2$ for the maximum path would yield the equation $\beta_{k,2}^{3/2} = 1$. Note that this exponent is different than the exponent calculated for the total path interpretation of time delay, which is different from the previous cases that minimize conduction time delay. We can rewrite this expression as

$$r_{k+1,1}^{3/2} = r_k^{3/2} \tag{A 38}$$

An analogous expression can be written for the maximum path length calculation, with $r_{k+1,2}^{3/2} = r_k^{3/2}$. Focusing on just the relationship between one daughter and the parent branch, this relationship is **area-decreasing**. However, given that the radius of the second daughter does not affect this relationship, is it possible that it is in fact area-preserving or increasing.



royalsocietypublishing.org/journal/rsif J R Soc Interface 0000000
